# Co-Metabolic Growth and Microbial Diversity: Keys for the depletion of the α, δ, β and γ-HCH isomers

**DOI:** 10.1101/2024.06.25.600594

**Authors:** Giacomo Bernabei, Giampiero De Simone, Simone Becarelli, Riccardo Di Mambro, Alessandro Gentini, Simona Di Gregorio

**Affiliations:** Depertment of Biology, University of Pisa, Italy; Teseco Bonifiche srl, Italy

**Keywords:** hexachlorocyclohexane, microbiomes, quantitative 16S DNA metabarcoding, α, δ, β, γ-HCH isomers, *Chryseobacterium* sp., *Asinibacterium* sp.

## Abstract

The objective of this study was to select and enhance microbiomes capable of degrading the α, δ, β and γ-HCH isomers. These microbiomes were isolated and enriched from an HCH-contaminated dumpsite in Italy, both in the presence of HCH isomers (1:1:1:1) as the sole carbon sources and under co-metabolic growth conditions in presence of glucose (0.1%). Four microbiomes were assessed for their relevant metabolic capabilities. A quantitative metabarcoding approach was employed to analyze the compositional evolution of the four microbiomes during the enrichment phase and the phase of tsting of the HCH isomers degradation kinetics. The use of a co-metabolic substrate during enrichment process was essential for selecting microbiomes with higher biodiversity. All microbiomes efficiently degraded the α, δ, and γ-HCH isomers. The highest efficiency in the β-HCH degradation capacity was associated to the highest biodiversity of the microbiome, and the involvement of *Chryseobacterium* and *Asinibacterium* sps. has been proposed for a recorded increment in bacterial load during the HCH degradation process.

**Statement of environmental implications:** Soil contaminated with hexachlorocyclohexane (HCH), including all four isomers, poses a significant risk to environmental and public health. This study isolates and selects microbiomes capable of degrading HCH, demonstrating their degradation efficiency using GC-MS analysis, and studies the microbial communities through metabarcoding of both the initial soils and the selected microbiomes. The contaminated soil originates from the historically polluted area of Italy known as SIN-Valle del Sacco. Developing and optimizing microbiome selection techniques for application on contaminated sites can significantly enhance soil bioremediation, thereby reducing contamination and protecting the environment.

## 1. Introduction

The technical 1,2,3,4,5,6-Hexachlorocyclohexane (t-HCH) or HCH is an organochlorine mixture of isomers^1^. A total of four stable isomers have been identified and quantified in HCH formulations, the α-(60 to 70%), β-(5 to 12%), γ-(10 to 12%), δ-(6 to 10%) and ε-(3 to 4%)^2^.

Among these isomers, only the γ-HCH, also known as Lindane, shows insecticidal properties, whose production justified the one of the HCH worldwide, for decades^3^. In several countries, Lindane has been used as a pure formulation (>99%)^4^, claiming for processes of purification, causing the dumping as wastes of the α, β and δ-isomers in large quantities, with consequent pollution of neighboring soils and aquifers^5^. All HCH isomers are acutely and chronically toxic^6^. Even though HCH production has been progressively worldwide dismissed, and restrictions have been imposed on its usage^7^, it is still of significant toxicological concern. The production of 1 ton of Lindane generated about 8-12 tons of different isomers as wastes. In addition to the widespread pollution sources, stemming from Lindane usage and production, significant challenges arise also from specific locations, where unused Lindane remains stockpiled globally. The concentrations of HCH in the environment near stockpiles and productions sites can reach exceptionally high levels, typically measured in grams per kilogram of soil or milligrams per liter of water^6^. Remediation of contaminated environmental matrices is needed, however, due to extension of the problem, a sustainable approach is mandatory. A bio-based approach to the problem is the most promising in terms of environmental and cost sustainability. With the aim to design effective processes of decontamination it must be mentioned that the complete microbial mineralization of lindane has been observed under aerobic conditions^8^. Microbial aerobic degradation of the HCH isomers has been observed frequently in both mixed and pure microbial cultures^9^. The complete aerobic degradation pathway of Lindane has been characterized in different bacterial taxa, predominantly from the family Sphingomonadaceae, where the pivotal role of the *lin* genes has been described^2,7^. However, the catabolic pathway for the major isomers other than Lindane has not been fully elucidated, even though the putative role of the *lin* genes has been suggested^2^. The β-HCH is the most recalcitrant of all the HCH isomers and does not undergo mineralization easily^9,10^. Its stability is due to its chemical structure, a fully equatorially substituted HCH isomer^11^. This appears to be a barrier to the dehalogenation reactions, which are the first steps in the degradation of HCH isomers^11^. In terms of equatorial substitution, the δ-HCH isomer has five of the chlorines in the critical position, also posing a barrier to the first attack on the molecule. The genes involved in the first dehalogenation of HCH isomers are harbored by the *lin* pathway and the *lin*A and *lin*B genes trigger the start of degradation, while other genes in the pathway facilitate subsequent degradation steps^12^. The α- and γ-HCH isomers are mineralized by the *lin* pathway under aerobic conditions by several taxa, while β- and δ-HCH isomers have been reported to be degraded only to certain hydroxylated metabolites by a few bacterial strains^12^. Dedicated and even interspecies diverse route of dehalogenation have been described in *Sphingobium* sp.. For the equatorial dehalogenation of the β-isomer, *Sphingobium* sp. was described as adopting both dehalogenation via the same mechanism as that of the axial chlorines in γ-HCH, and, for the equatorial chlorine, the same mechanism as that of in β-HCH^12,13^.

Although multiple bacterial strains have been described as capable of partially transforming more than one HCH isomer, an unequivocal description of single strains efficiently mineralizing all the HCH isomers at once is still missing. This evidence might be the actual bottleneck for the design of a bio-based approach to HCH decontamination in the environment.

To reach the aim of a transformation of all the HCH isomers in bio-based approach for contaminated environmental matrices, the recruitment of competent bacterial consortia might be beneficial, as it is well known that microbiomes work better than single strains, with reference to the degradation of recalcitrant organic compounds^14–16^. In the case of HCH a microbial cooperative process of degradation of all the HCH isomers is looked for. Microbial consortia capable of degrading multiple HCH isomers at once have already been isolated^17,18^, however, little is known about their composition. In this context, it should be mentioned that in the case of a bio-based approach to the decontamination of environmental matrices, bioaugmentation of selected strains capable of depleting contaminants, is usually considered as a promising approach. However, the success of bioaugmentation can significantly depend on the identification and isolation of suitable microbial degrading strains, as well as their ability to survive and remain active once introduced into the intended environment. In this context, microbiomes autochthonous to the matrices are mandatory to trigger degradation processes in the ecological habitat of origin. Moreover, they can establish syntrophic relationships leading to the decomposition of recalcitrant contaminants, when the individual degradation is unattainable, making these intricate community interactions crucial for depleting contaminants from environmental matrices. Grasping the cooperative function, communication, and actions of these communities in bioremediating the environment, holds significant interest and importance.

Thus, the scope of the present work was the selection and enrichment of microbiomes capable of depleting all the four isomers of HCH. The microbiomes were isolated and enriched from a HCH contaminated dumpsite in Italy, both in presence of HCH as the sole carbon source and in co-metabolic growth conditions. Four microbiomes were characterized for the metabolic capability of interest. A quantitative metabarcoding approach has been adopted to taxonomically characterize the compositional evolution of the four promising microbiomes, whose HCH kinetics of degradation were described, with a particular focus on the δ-HCH and the β–HCH isomers. The involvement of a co-metabolic substrate in the isolation and enrichment of HCH degrading microbiomes resulted to be mandatory to select microbiota with a higher biodiversity. All the microbiomes depleted efficiently the α, δ, γ–HCH isomers. The highest microbiome biodiversity was correlated to the highest efficiency in the β–HCH isomer depletion with the putative involvement of *Chryseobacterium* and *Asinibacterium* sps., showing an increment in bacterial load during the β–HCH isomer depletion phase.

## 2. Material and Methods

### 2.1 Chemicals and contaminated soils

Two soil samples (AN7-8 and AN10-11) were collected at Colleferro Site (41.747753, 12.978449). The contamination levels are reported in Table 1.

**Table 1.**
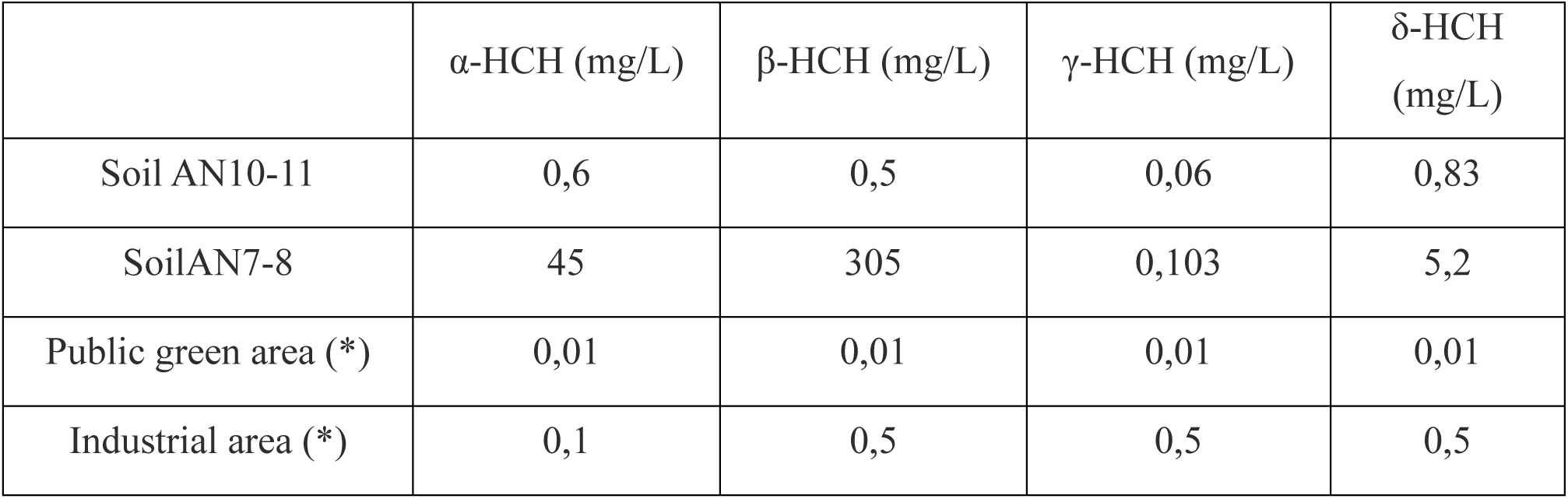
The table displays the concentrations in mg/L of the 4 detected HCH isomers in the soil samples: α, β, γ and δ. (*) Limits of allowed concentration in Italy based on soil area exploitation

All analytical grade reagents were purchased from Merk Life Science S.r.l. (Milano). The HCH contains isomers α, β, γ and δ in 1:1:1:1 ratio (HCH, PESTANAL®, analytical standard, mixture of isomers α:β:γ:δ=1:1:1:1).

### 2.2 Enrichment of HCH degrading microbiota

The HCH degrading microbiota were isolated by two enrichment phases, the adaptation and the selection phase, in MSM (Mineral Salts Medium) supplemented with increasing concentration of technical HCH (HCH). MSM derives from a modified version of the medium used to isolate *Xanthomonas* sp. ICH12^19^ and contains (per L of medium): KH2PO4, 170 mg; Na2HPO4, 980 mg; (NH4)2SO4, 100 mg; MgSO4, 4.87 mg; FeSO4, 0.05 mg; CaCO3, 0.2 mg; ZnSO4, 0.08 mg; CuSO4·5H2O, 0.016 mg; Na2MoO4·2H2O, 0.05 mg; MnSO4, 0.05 mg; H3BO3, 0.006 mg.

In relation to the adaptation phase, soil samples were incubated in MSM or in MSM amended with glucose 0.1% v/v (MSMglu) for 14 days. The microbiota deriving from the enrichments in presence of glucose were labeled as positive (Pos: AN7-8Pos and AN10-11Pos), the one deriving from enrichment in absence of glucose were labeled as negative (Neg: AN7-8Neg and AN10-11Neg). In details, 15 g of deeply homogenized AN7-8 and AN10-11 soil samples were incubated in 250 mL Erlenmeyer flasks containing 135 mL of sterile MSM and incubated on a rotary shaker at 100 rpm at 25 ± 1°C for 14 days in the dark (AN7-8Neg and AN10-11Neg). In parallel, different aliquots (15 g each) of the same deeply homogenized soil samples were incubated in the same conditions but in MSMglu (AN7-8Pos and AN10-11Pos). At the 15th day, the flasks were left undisturbed for 30 minutes for the settling of soil particles. Successively, 100 mL of the supernatant consisting in the spent growth medium was removed and replaced with an equal volume of fresh MSM and MSMglu amended with 10 mg/L of HCH. The incubation lasted in the same condition until day 42 (phase of adaptation of the soil microbiome to the new growth conditions in presence of HCH).

The selection phase of the HCH degrading microbiota started at day 43, during which the HCH concentration increased up to 100 mg/L. In details, the flasks were deeply vortexed and 50 mL from each flask were collected and used as inoculum for new flasks containing 100 mL of sterile MSM and MSMglu initially amended with HCH 10 mg/L. At day 72, 100 mL of spent growth medium was removed from each bottle and replaced with an equal volume of fresh MSM and MSMglu media amended with 10 mg/L of HCH. The process was repeated for two times up to day 135, time of incubation when, following the same procedure, the concentration of HCH in the MSM and MSMglu was increased up to 50 mg/L. The same procedure was adopted also at day 163 when the HCH concentration was increased at 100 mg/L. Both at day 72, 135 and 163 three biological samples of the total volume of 100 uL were collected for the extraction of the metagenomic DNA.

### 2.3 Microbiome HCH degradation testing phase

The testing phase was dedicated to measuring the HCH depletion capacity of the microbiomes obtained by the four enrichment conditions. The test was conducted in presence of 100 mg/L HCH for 20 days of incubation. At day 164 a total of 87 sterile tight closed glass vials, volume of 20 ml, were prepared as follows: 10 mL of MSM, MSMglu were inoculated with 100 uL of bacterial cell suspension collected by vortexed enrichment flasks (AN7-8Neg and Pos; AN10-11Neg and Pos). Three vials per time of analysis (0, 3, 7, 10, 20 days) were prepared. Three non-inoculated vials were prepared as control to check the abiotic loss of HCH. Three vials for each microbiome (for a total of 12 vials) were prepared and incubated as previously described to collect the cell pellet for the extraction of the metagenomic DNA after 20 days of incubation of the different microbiomes in presence of 100 mg/L HCH. The cell supernatants were extracted and analyzed for HCH content. All the glass vials were incubated on a rotary shaker at 100 rpm at 25 ± 1°C in the dark. At the end of each incubation time three vials for each experimental condition for a total of 75 vials were extracted with n-hexane and analyzed for HCH content by GC-FID. At day 20, the 12 vials extracted for bacterial metagonomic DNA, pelleting the cell suspension by centrifugation (8000 rpm for 20 min) were also analyzed for HCH content.

### 2.4 HCH quantification by GC-FID

The entire volume of the testing phase vials was extracted in an equal volume of n-hexane containing 24 ppm of fluorene as an internal standard by vortexing for 2 minutes. Subsequently, a portion of the non-polar fraction was collected and treated with anhydrous Na2SO4, previously dried in an oven at 105°C for 90 minutes, to remove the water dissolved in the extraction solvent. A volume of 1 µL of the extract was injected into an Agilent 7890B gas chromatograph (GC), equipped with the Flame Ionization Detector (FID), and a column with a length of 20 m and an internal diameter of 0.18 mm, internally coated with a 0.18 µm film composed of a non-polar stationary phase containing 5% diphenyl-polysiloxane (HP5-MS, Agilent). The injection was performed in split-less mode at 280°C: Constant temperature injector at 280°C, septum purge flow rate 3 mL/min, injector purge flow rate 40 mL/min after 1 minute of split-less mode; carrier gas (helium) isocratic flow rate 0.8 mL/min. Oven temperature programming: 80°C for 2 minutes, ramp to 100°C at 20°C/min, held for 1 minute, ramp of 27.5°C/min to 240°C, immediately followed by a ramp to 250°C at 14°C/min, held for 4 minutes. Total run time of 14 minutes. External calibration was performed with HCH standards containing fluorene as an internal standard.

### 2.5 Quantitative 16S rDNA metabarcoding

The total DNA were extracted from 500 mg of soil samples or 100 mg of bacterial cell pellet using a FastPrep 24™ homogenizer and FAST DNA Spin kit for soil (MP Biomedicals, California, USA). The cell pellets were collected after 72, 134, 163, 184 days during the microbiome enrichment and testing phases. The quantity of DNA was measured using a Qubit 3.0 Fluorometer (ThermoFisher Scientific, Milan, Italy). The DNA purity and quality was determined spectrophotometrically (Biotek Powerwave Xs Microplate Spectrophotometer, Milan, Italy) by measuring absorbance at 260/280 and 260/230 nm.

From both soil and enrichment samples a total of 200 ng of DNA was utilized to produce paired-end libraries and for sequencing the V4–V5 hypervariable regions of the bacterial 16S rRNA gene by using as primers, 515F forward primer (5′-GTGCCAGCMGCCGCGGTAA-3′) and 907R reverse primer (5′-CCGTCAATTCCTTTGAGTTT-3′). The libraries for Illumina sequencing were prepared by Novogene (Novogene Company Limited Rm.19C, Lockhart Ctr., 301–307, Lockhart Rd. Wan Chai, Hong Kong) using the Illumina NovaSeq 6000 platform. NCBI sequence access code: PRJNA1114457.

The paired-end reads resulting from the sequencing of the 16S libraries of various metagenomes were assigned to samples based on a unique barcode, which was subsequently removed^20^ (Cutadapt v 4.6). Subsequently, the forward and reverse reads were assembled, filtered for read quality (minimum Phred-scaled quality score = 10), cleaned from potential chimera sequences, and assigned to clusters represented by each Amplicon Sequence Variant (ASV) using the DADA2 plugin v 1.26^21^ of Qiime2 v 2023.2.^22,23^. The annotation of ASVs produced by DADA2 was performed using the RESCRIPt plugin v 2023.2.0^24^ in Qiime2 with a classifier trained on the respective hypervariable region extracted from the Silva 138 99% 16S sequences database. To quantify the microbial load (number of normalized ASVs / mL of MSM) for each microbiota, four exogenous DNA spikes at known concentrations were included in the DNA samples before sequencing, as suggested by Harrison et al.^25^.

Microbial load was calculated as follow:

> 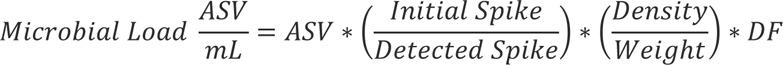
>
> Equation 1. Microbial load is calculated as number of ASV per mL of MSM. ASV is the number of ASV calculated as reported; Initial Spike is the quantity of synthetic DNA put in the sample before sequencing; Detected Spike is the detected quantity of the synthetic DNA after sequencing; Density and Weight were measured for the initial samples used for DNA extraction; DF refers to the Dilution Factor used to obtain the standardized concentration of 200 ng/mL of DNA required for the library preparation step.

The statistical significance of the microbial load was assessed using mixed-effects linear regression, starting from log-normalized counts, as suggested by Anderson et al.; followed by paired T-tests as post-hoc tests^26^ (lme4 v 1.1-33, rstatix v 0.7.2).

Differential abundance analysis of microbial taxa was carried out using ANCOMBC 2.0 and ggplot2 v 3.4.2 and VennDiagram (v1.7.3) was used for creating graphs^27,28^.

The abundance of ASVs was normalized using the coverage method (0.9950 ± 0.0025). Alpha-diversity was assessed by calculating Hill numbers, corresponding to the inverse of the Simpson index (Hill-Simpson) and for Hill-Shannon, along with the Chao1 index (metagMisc_R v 0.5.0)^29–31^. The statistical significance of these indices was evaluated through non-parametric repeated measures ANOVA, followed by post-hoc Dunn tests^32^ (nparLD v 2.2, rstatix v 0.7.2). Additionally, rarefaction curves of observed species were generated based on the coverage reference. To estimate beta-diversity, Principal Coordinate Analysis (PCoA) based on weighted UniFrac distances was performed.^33,34^ (Phyloseq v 1.41.1, Vegan v2.6-4).

The core microbiome was identified by selecting taxa that had an abundance exceeding 0.1% within each sample and were present in at least 50% of the samples. More precisely, these taxa needed to be detected in at least two out of the three sequencing replicates per sample^23^ (microbiome v 1.24.0).

## 3. Results

### 3.1 Soils Alpha Diversity

The biodiversity indexes of AN7-8 and AN10-11 soil samples, from which the HCH microbiomes derived, are shown in Figure 1. The Chao1 estimator and Hill-Shannon indexes showed a higher richness in bacterial taxa in AN7-8 soil sample with reference to the AN10-11 one. The differences recorded are significant only in the case of the Chao1 index, however the Hill-Shannon and Hill-Simson indexes showed tendencies worth to be considered. Although both Chao1 and Hill-Shannon diversity indices indicate higher levels of alpha diversity for AN7-8, the Hill-Simpson index shows an opposite trend. This discrepancy suggested a higher abundance of rare and less represented bacterial species in the AN10-11 soil sample compared to the AN7-8 one.

**Figure 1.**
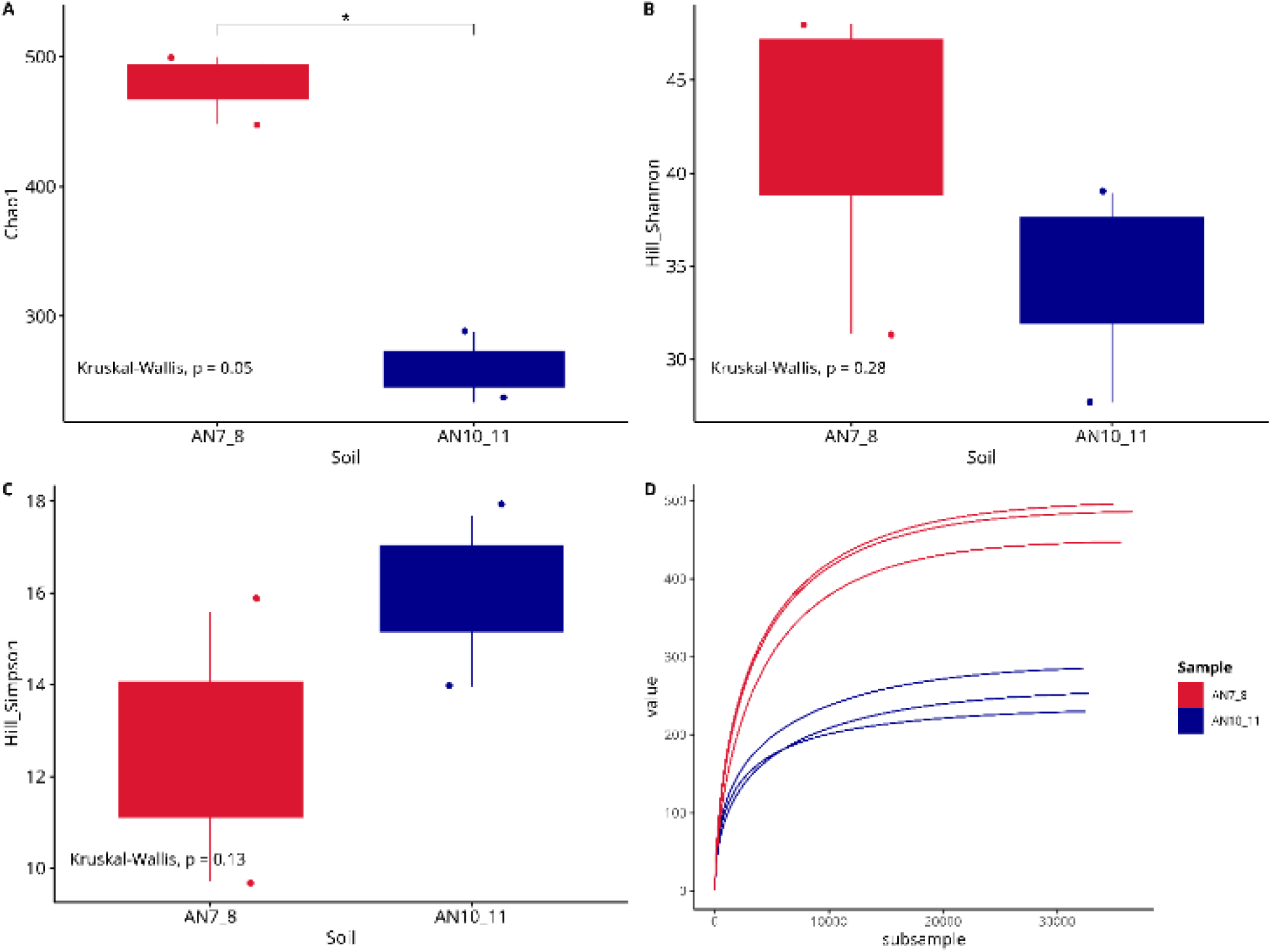
α diversity of soil samples AN7-8 and AN10-11. Chao1 index (A) Hill–Shannon (B) Hill–Simpson index (C), each calculated after rarefaction to a coverage of 99.5%, and rarefaction curves of observed species of bacterial community composition (D). Box and whiskers represent the minimum (Q0), 1st quartile (Q1), median (Q2), 3rd quartile (Q3), and maximum (Q4) of each group. Reported p-value is calculated by the Kruskal–Wallis test (α = 0.05). Post hoc statistical test is based on the Dunn test with Benjamini–Hochberg correction for multiple comparisons.

### 3.2 HCH depletion during the testing phase of the microbiota

Four microbiomes were obtained at the end of the selection phase after 163 days of incubation. Two microbiomes were obtained by the exploitation of HCH as sole carbon source, the AN7-8Neg and the AN10-11Neg. Two microbiomes were obtained with glucose as co-substrate with HCH, the AN7-8Pos and the AN10-11Pos. The 4 microbiomes derived from a phase of adaptation of 15 days in absence of HCH, were successively incubated in the presence of gradually increasing concentrations of HCH, from 10 to 100 mg/L. The capacity of the four microbiomes to deplete this last concentration in 20 days of incubation was tested. Results obtained with reference to the microbiome capacity to deplete 100 mg/L HCH are shown in Figure 2.

**Figure 2.**
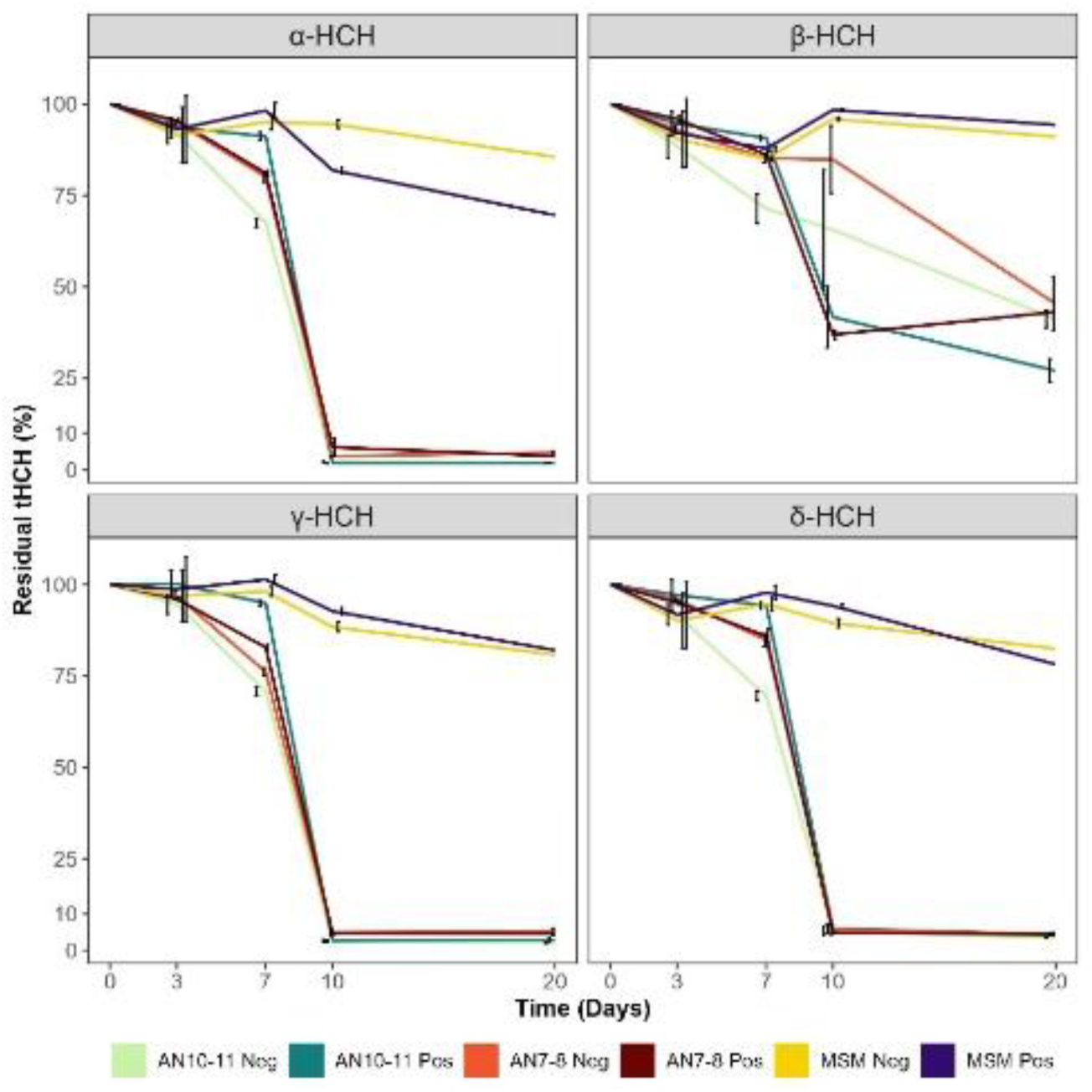
The residual t-HCH isomers are represented with reference to days of incubation of the 4 microbiomes and 2 control sample (MSM Neg, MSM Pos) in presence of 100 mg/L t-HCH.

The four different microbiomes, after 10 days of incubation at 100 mg/L HCH, showed a depletion of 95% for the α-, γ-, and δ - HCH isomers. In relation to the β isomer the AN7-8Pos, AN7-8Neg, and AN10-11Neg microbiota reached 50% of depletion after 20 days of incubation. On the other hand, during the same time interval, the AN10-11Pos microbiota reached up to 75% of depletion of the β isomer. Also notable are the trends of abatement. In presence of glucose (MSMglu, AN7-8Pos and AN10-11Pos) the kinetics of depletion of the β isomer were faster, reaching the 50% of depletion after only 10 days with reference to the enrichment incubated in absence of glucose (MSM, AN7-8Neg and AN10-11Neg), showing, after the same time interval, the 25% of depletion. The abiotic loss of HCH, in the adopted testing phase conditions, accounted for values lower than 25%.

### 3.3 Bacterial community diversity

A metabarcoding approach was adopted to compare the bacterial diversity of the soil samples utilized to enrich the 4 HCH degrading microbiomes and these latter. Figure 3 shows the β-diversity with reference to the comparison in biodiversity of the bacterial communities characterizing the four HCH depleting microbiomes during their enrichment and testing phase and the two soil samples the 4 microbiomes derived from.

**Figure 3.**
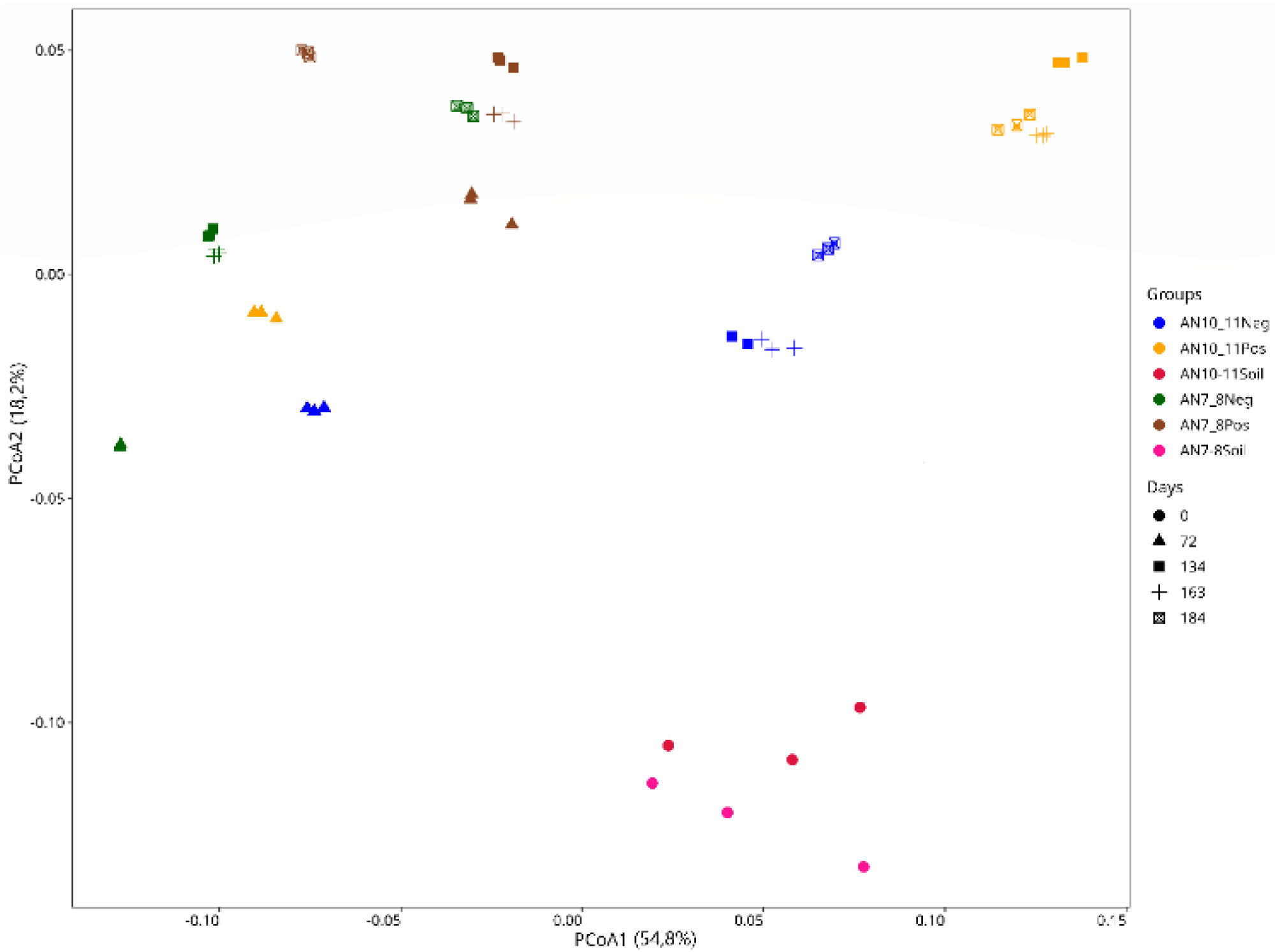
Principal Coordinate Analysis (PCoA) of bacterial communities, conducted based on weighted UniFrac distances for each triplicate per sample. Point colors represent the analyzed samples, while shapes indicate the temporal category. The amount of total variance represented by

The analysis was performed to highlight potential similarities and differences in the overall taxonomic composition of the evolving microbiomes. The first time point of the analysis for the microbial ecology of the HCH depleting microbiomes was selected at day 72 to analyze microbial communities reasonably significantly affected by the incubation in presence of an excess of HCH, more precisely incubated for 60 days at 10 mg/L of HCH, after 15 days of adaptation in absence of HCH. The microbial diversity was analyzed also at day 134 just before the increase of HCH concentration at 50 mg/L (119 days at 10 mg/L HCH) and at day 163 just before the increase of HCH concentration at 100 mg/L (29 days at 50 mg/L HCH). The microbial diversity of the different HCH degrading microbiomes was analyzed at the end of the testing phase (day184).

The overall variability analyzed by PCoA, represented by the sum of the weights of each axis (PCoA1 and PCoA2), accounted for 73% of the total diversity. Within the two reported dimensions, a clear segregation was observed between the soil microbial communities and the one of the evolving microbiomes during their phases of enrichment and testing. Based on PCoA1, the former remains on the lower side of the graph, while the latter occupies the upper side. On the other hand, the distance between the bacterial diversity of the HCH depleting microbiomes at day 72 of the enrichment phase, in presence of 10 mg/L HCH and the one of the microbiomes at the remaining time of analysis was higher with reference to the distance of the bacterial diversity of the microbiomes evolving at higher HCH concentrations and the one of the HCH degrading microbiomes. The effect is particularly evident for microbiome AN10-11Pos.

### 3.4 Quantitative 16S rDNA metabarcoding

A quantitative metabarcoding approach was adopted to analyze the evolution of the bacterial diversity of the 4 HCH degrading microbiomes during the enrichment and testing phases. Figure 4 shows the evolution of the most abundant bacterial taxa (≥ 1% in relative abundance), expressed as absolute bacterial counts.

**Figure 4.**
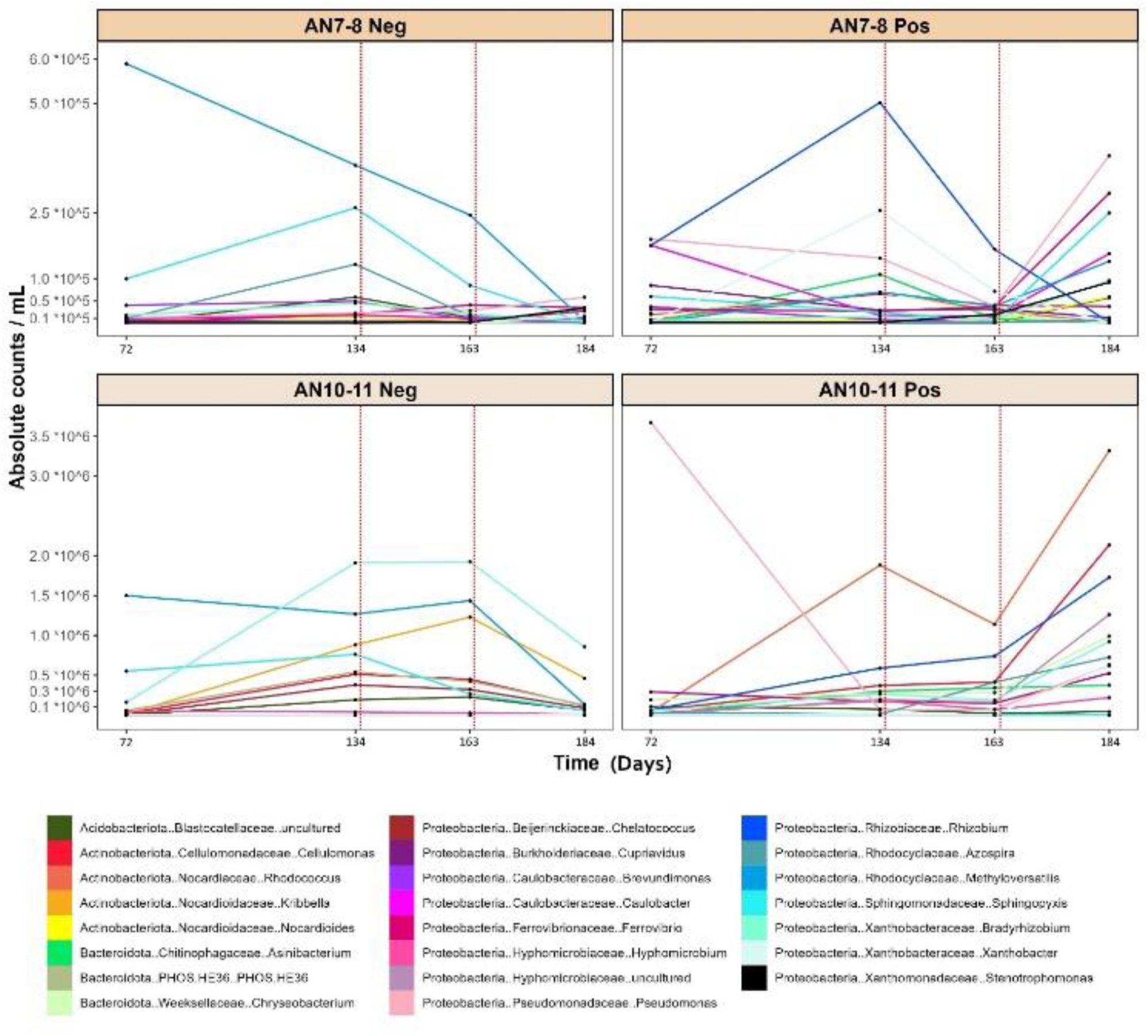
The absolute bacterial counts expressed as absolute counts per milliliter of growth medium of the 4 microbiomes during the enrichment and testing phase. Each color corresponds to a specific bacterial taxon. Vertical dotted lines at day 134 and day 163 correspond to the increase of tHCH in the medium, from 10 mg/mL to 50 mg/mL and 100 mg/mL, respectively.

An arbitrary threshold in total counts was adopted to represent the trend of the most abundant species and those reasonably involved in the process. Results obtained show that increasing the concentration of HCH from 10 to 50 mg/L (day 134) and from 50 to 100 mg/L (day 163), had either a bacteriostatic or bactericidal effect on most of the observed taxa of the microbiomes incubated in absence of glucose (AN7-8Neg and AN10-11Neg). The effect was evident also during the testing phase between day 163 and 184. On the other hand, the absolute counts of most of the observed taxa significantly increased during the testing phase (day 163-184) for the microbiomes incubated in presence of glucose (AN7-8Pos and AN10-11Pos). Interestingly, the observed increase in AN7-8Pos affects almost exclusively taxa belonging to the phylum Proteobacteria, while for AN10-11Pos the taxa that showed an increase in their absolute abundance are distributed among the phyla Acidobacteriota, Actinobacteriota, Bacteroidota and Proteobacteria.

In the microbiota AN7-8Neg, the bacterial taxa *Methyloversatilis, Sphingopixis* and *Azospira*, all belonging to the phylum Proteobacteria, were the most abundant at day 134 and 163. Their abundance decreased after day 163 during the testing phase.

In the AN7-8Pos microbiome, the bacterial taxa *Rhizobium, Xanthobacter, Pseudomonas* and *Ferrovibrio*, all belonging to the phylum Proteobacteria, were the most abundant taxa at day 134. *Rhizobium* and *Xanthobacter* increased their abundance at day 163, along with the Bacteroidota *Asinibacterium*. Meanwhile, *Ferrovibrio* is linked to a decrease in abundance and *Pseudomonas* remains unchanged. After the increase of HCH concentration at day 134, the abundance of all the aforementioned taxa decreased. Interestingly, from day 163 to day 184, the abundance of multiple bacterial taxa increased significantly; these taxa are *Pseudomonas*, *Ferrovibrio*, *Sphingopyxis*, *Caulobacter*, *Methyloversatilis*, *Bradyrhizobium* and *Stenotropomonas*, all belonging to the Phylum Proteobacteria.

In the AN10-11Neg microbiome, the Proteobacteria *Bradyrhizobium*, *Methyloversatilis* and *Sphingopyxis* are the most abundant taxa at day 72 and they either increased or maintained their abundance at day 163. The bacterial taxa *Kribbella* and *Cellulomonas*, belonging to the Phylum Actinobacteriota, along with the Bacteroidota *PHOS.HE36*, also increased their abundance significantly from day 72 to day 134. After the increase of HCH concentration, they overall maintained a stable abundance until day 163, which then decreased at day 184.

In the AN10-11Pos microbiome, the taxon *Pseudomonas* is the most abundant at day 72, although its abundance decreases drastically at day 134. On the other hand, the Actinobacteriota *Kribbella* and the Proteobacterium *Rhyzobium* increased their abundance at day 134. Successively, *Kribbella* decreased in abundance up to day 163. Similarly to AN7-8Pos microbiome, also in the AN10-11Pos, the abundance of almost all the selected taxa increased from day 163 to day 184. However, the increase was not restricted to the Phylum Proteobacteria, more precisely with *Rhyzobium*, *Sphingopyxis*, *Azospira*, *Xanthobacter* and *Ferrovibrio* sps., but involving also the Actinobacteriota *Kribbella* and *Cellulomonas*, along with the Bacteroidota *Chryseobacterium*.

### 3.5 Core microbiome analysis

To infer putative function to bacterial taxa exclusive to AN10-11Pos microbiome, depleting the HCH isomers with a higher efficiency, with reference to the other microbiomes, a core microbiome analysis was adopted. The core microbiomes of the four HCH depleting microbiomes (AN7-8Pos, AN7-8Neg, AN10-11Pos, AN10-11Neg) at the end of the testing phase and the one of the soils of origin were compared and represented with the Venn diagram (Figure 5). The bacterial strains recovered over the threshold selected to define the core microbiome and exclusively recovered in AN10-11Pos, belonged to Proteobacteria, Acidobacteriota, Actinobacteriota and Bacteroidota and are reported in Table 2. Two of them were not characterized at genus level. The *Asinibacterium* and the *Chryseobacterium* sps. were represented by two strains respectively. On the other hand, three strains of the Hyphomicrobiaceae family, recovered as defining the core microbiomes were present both in the AN10-11Pos microbiome and in the soil of origin.

**Figure 5.**
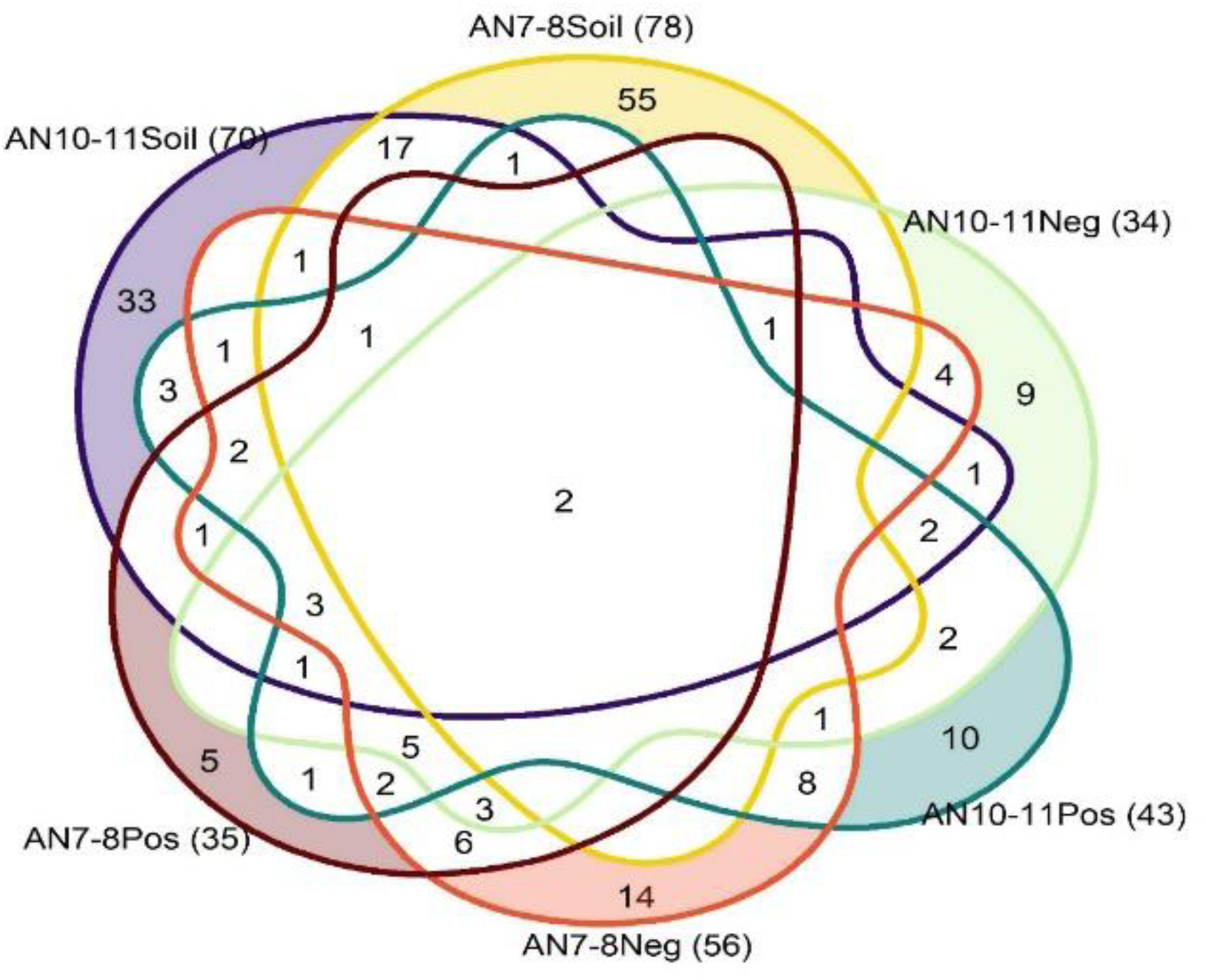
This Venn diagram, constructed using the R package VennDiagram (v 1.7.3), depicts the shared taxa among microbial communities derived from six samples representing various microbial habitats: AN10-11 (both Positive and Negative at the 184th day), AN10-11 soil, AN7-8 (both Positive and Negative at the 184th day), and AN7-8 soil. The core microbiome presented in the diagram was determined by identifying taxa with an abundance exceeding 0.1% within each sample and with a prevalence of at least 50%. Specifically, taxa were required to be present in at least two of the triplicates of sequencing per sample. This methodology aims to highlight the taxa consistently present across replicate samples, providing insights into the stable microbial communities within the studied environments. The colored arrows indicate the taxa shared between AN10-11 Soil and AN10-11 Pos microbiome (blue arrow) and the taxa exclusive of the AN10-11 Pos microbiome (green arrow).

**Table 2.** The table lists the 3 taxa shared between AN10-11 soil and AN10-11Pos microbiome, as well as the 10 taxa exclusive to the AN10-11Pos microbiome at the end of the testing phase. For each taxon, the phylum, family, and genus are provided. These taxa correspond to those depicted in the Venn diagram shown in Figure 6. The colored arrows refer to taxa shown in Figure 5.

|  | Phylum | Family | Genus |
| --- | --- | --- | --- |
| → Taxa shared between AN10-11 soil and AN10-11Pos microbiome at the end of the testing phase | Proteobacteria | Hyphomicrobiaceae | *uncultured* |
|  | Proteobacteria | Hyphomicrobiaceae | *uncultured* |
|  | Actinobacteriota | Nocardioidaceae | *Nocardioides* |
| → Taxa present only in AN10-11Pos microbiome at the end of the testing phase | Proteobacteria | Dongiaceae | *Dongia* |
|  | Proteobacteria | Sphingomonadaceae | *Sphingomonas* |
|  | Acidobacteriota | Blastocatellaceae | *Stenotrophobacter* |
|  | Actinobacteriota | Nocardiaceae | *Nocardia* |
|  | Actinobacteriota | Ilumatobacteraceae | *CL500-29_marine_group* |
|  | Chloroflexi | JG30-KF-CM45 | *JG30-KF-CM45* |
|  | Bacteroidota | Chitinophagales | *Asinibacterium sp.1* |
|  | Bacteroidota | Chitinophagales | *Asinibacterium sp. 2* |
|  | Bacteroidota | Weeksellaceae | *Chryseobacterium sp.1* |
|  | Bacteroidota | Weeksellaceae | *Chryseobacterium sp.2* |

## 4. Discussion

The catabolic abilities of bacteria towards contaminants in the environment led to the design of bio-based processes aimed at harnessing bacterial metabolism for detoxifying contaminated matrices. In the case of HCH, the presence of various HCH isomers, coupled with the frequent inability of bacterial dechlorinases in the *Lin* pathway to attack all the four HCH isomers, makes the use of multi-species microbial consortia interesting within bioaugmentation approach for the decontamination of HCH contaminated matrices. In fact, microbial biodiversity is accompanied by broad gene pools and metabolic flexibility that amplify the catabolic capabilities of a microbial community with reference to the single strain. Despite microbiome importance, limited attempts are reported to depict how the mechanism of isolating such microbiomes influences their composition and their biotechnological potential, even though the importance of developing protocols to isolate microbiomes with high diversity and possible broad metabolic potential is evident.

The here adopted strategy to isolate HCH transforming microbiomes was based both on the concept that the microbiomes should use HCH as sole carbon source, and on the extensively described co-metabolic HCH degradation^35^. Both concepts were adopted and the effect of a co-metabolic carbon source on the microbiome biodiversity, and the eventual catabolic flexibility towards all HCH isomers, has been observed. The metabolic processes associated to the HCH transformation is accompanied by a very low production of reductive power for the bacterial cells, Consequently they are gaining advantage in recovering an alternative carbon source to transform the toxic HCH isomers. The here adopted strategy comprised also glucose as carbon co-substrate, for improving the capacity to enrich and characterize microbiomes with a high transformation efficiency of all the toxic HCH isomers. The enrichment strategy comprised also the selection of two different soils contaminated with diverse HCH concentrations, presenting a diverse bacterial ecology. The lower HCH level of soil contamination was associated to a lower bacterial biodiversity and a higher evenness within the colonizing bacterial species, with reference to the bacterial ecology of the soil characterized by a higher level of HCH contamination. The four obtained microbiomes were all efficient in the depletion of HCH, but the co-metabolic strategy of enrichment increased the capacity of the co-metabolically enriched microbiome to transform the β isomer. Moreover, the four microbiomes depleted the δ isomer with kinetics very similar to the one observed for the less recalcitrant α and γ isomers. To the best of our knowledge this is the first report where the recalcitrance to biodegradation of the α, γ and δ isomers resulted to be similar. In is worth mentioning also that the strategy of enrichment of the four microbiomes consisted in a long phase of incubation of the microbiomes to a relatively low HCH concentrations. The microbial communities changed in composition during the enrichment phase, acquiring a compositional resilience to the increment of the HCH concentration, as shown by the β-diversity analysis, after the long-lasting acclimatation at the lowest HCH concentration. The effect is particularly evident for the AN10-11Pos microbiome. Since the microbiome was deriving from the soil with a higher evenness among the bacterial taxa characterizing the soil matrix, it is reasonable to suggest that a long acclimatation phase might be favorable also for the rare species whose presence is assessed by the higher value of the Hill-Simpson index in the AN 10-11 soil. These latter might be rare bacterial taxa, capable to deplete the most recalcitrant β isomer, and favored by a co-metabolic growth condition to recover sufficient reducing power by chemical structure recalcitrant to the oxidation of the carbon-carbon bonds. Biodiversity and evenness are consistently responsible for the improved capacity of the selected microbiomes to deplete with a significant efficiency the β isomer. To confirm the assessment, it is worth mentioning that the quantitative metabarcoding showed that the microbiome composition varies depending on the soil of origin. Interestingly, the less contaminated soil characterized by a lower bacterial richness (Chao1 index) was the source of microbiomes (AN10-11Pos, AN10-11Neg) where the composing dominant bacterial species, during the enrichment and testing phase, were distributed among Actinobacteriota, Bacteroidota and Proteobacteria. On the other hand, the microbiomes deriving from the soil with the highest level of contamination and higher bacterial richness (AN7-8Pos, AN7-8Neg), showed dominant bacterial taxa all limited to Proteobacteria. It is reasonable to assume that he initial evenness of the soil of origin, higher for AN10-11 (Hill-Simpson), might have influenced the evenness of the resulting microbiomes. However, nor the evenness and the richness of the microbiomes was relevant for the transformation of the HCH isomers, except for the β one. On the other hand, the highest efficiency of depletion for the β isomer was associated to the co-metabolic consumption of glucose, that determined the blooming of a bacterial taxa belonging to Proteobacteria but also Actinobacteriota, Acidobacteria and Bacteroidota. The blooming of bacterial taxa all belonging to the Proteobacteria (AN7-8Pos) resulted to be less efficient in the β isomer depletion.

The β isomer has been reported as transformed by *Sphingomonas paucimobilis*^36^ and several strains of Pseudomonadales (e.g., *Pseudomonas aeruginosa*^37^, *Burkholderia*^38^, *Rhodococcus*^39^, and *Mycobacterium*^40^ sps.). The reported data suggest that the metabolic capacity of β isomer transformation is distributed among different bacterial families. Our results confirmed the assessment and suggested that an enrichment co-metabolic approach implement the microbial biodiversity and it is mandatory for improving the catabolic capacity of the enriched microbiomes.

The quantitative metabarcoding is a rigorous instrument of interpretation of the response of the single bacterial genera to the changing environment and, in this case, to the increment in HCH concentration. The enrichment approach was associated to the blooming od dominant genera already described as involved in HCH transformation. More precisely the *Pseudomonas*, *Cupriavidus*, *Chriseobacterium* and *Rhodococcus* sps., were all already described as bacterial genera harboring species that are characterized in literature as HCH and/or OCPs (organochlorine pesticides) degraders^39,41–44^.

*Nocardioides* sp. was also described as one of the dominant genera characterizing the microbiomes, and in literature, the strain PD653 has been described as capable of the β-HCH dehalogenation^45^. Although the experimental conditions are not directly comparable, the referenced study reported that strain PD653 could degrade 5.7 ppm of β-HCH in 48 hours. This performance could be evaluated as suboptimal compared to our, since we achieved a 55% reduction of the same isomer in 10 days, however, starting from a concentration of 100 ppm of all four isomers. Moreover, the adopted approach resulted in the selection of microbiomes highly specialized in depleting all the HCH isomers, including the most recalcitrant δ and β isomers, albeit with different kinetics. Results were encouraging in this direction when compared to other case studies^9,46^, reporting an average depletion rate of 0.026 µg/h, whereas the here reported rates were approximately 2.08 µg/h for the α, γ, and δ isomers, and 0.93 µg/h for the β isomer. On average, our microbial consortia were 50 to 100 times faster at degrading the four isomers of t-HCH.

*Xanthobacter* sp. also dominant in this experimentation, has been described in literature as harboring a *Lin*B gene encoding for HCH dehalogenation^47^, and even though not yet directly involved in HCH dehalogenation, the combination of our data and the one of the other authors suggest the involvement of the genus in HCH depletion. Despite not being directly characterized as HCH transforming bacterial species, candidates belonging to the genera *Brevundimonas*, *Cellulomonas* and *Hyphomicrobium* were found to be part of microbial consortia capable to deplete HCH^48–50^. Similarly, a *Stenotrophomonas* sp. strain has been reported to grow on minimal medium with γ-HCH as sole carbon source, and *Caulobacter vibrioides* has shown to bioaccumulate the pollutant^51,52^.

Taxa belonging to the genera *Rhizobium*, *Bradirhizobium*, *Sphingopyxis* and *Kribbella* were found in the microbial communities of dumpsites, agricultural soils and pond sediments polluted by HCH^53–55^. Moreover, in an experiment regarding the effects of organochlorides in the microbial diversity of the roots of *Phragmites australis*, the *Methyloversatilis* sp. was found to be susceptible to the presence of a cocktail of pollutants containing γ-HCH, as its relative abundance decreased in the polluted samples when compared to the control^56^.

Finally, despite the fact that they are not described for HCH transformation, the genera *Azospira*, *Chelatococcus*, *Ferrovibrio* and *Asinibacterium* are described for dehalogenation capacities^55,57^.

It is worth mentioning that the blooming of already described HCH transforming taxa at the increase of HCH concentration was evidently favored by the presence of a co-metabolic substrate, suggesting that even though reducing power for microbiome propagation was sufficient up to 50 mg/L of HCH as a sole carbon source, the increment to 100 mg/L was accompanied to a decrease in all the taxa in absence of glucose. The insufficient production of reducing power due to the paucity of energy content of the halogenated isomers might be the reason of the microbial load loss. However, the accumulation of toxic intermediates of degradation cannot be excluded. The annotation of the metagenome of the microbiomes is mandatory to depict functional features involved in the evolution of the catabolic capacities of the microbiomes grown in different metabolic conditions. However, to assume a specific role for a specific taxon in the observed kinetics of HCH depletion, the strains exclusive to the AN10-11Pos, that might be associated to the highest efficiency in the corresponding β isomer depletion, were recovered by the core microbiome analysis. Most of the observed taxa were under the selected threshold of representation of the microbial load. Only the *Chryseobacterium* and *Asinibacterium* sps., that in the microbial load is aggregated to genus species, increased in the testing phase in presence of glucose with counts significant high to be represented by the microbial load. *Chryseobacterium* sp. has been reported as involved in the transformation of organochlorine pesticides^44^ and forming biofilm on biochar exploited for enhancing the natural attenuation of organochlorine pesticides^58^. A specific intervention of the taxa in determining the highest efficiency in the β isomer depletion cannot be excluded, as well as a synergistic intervention of the remaining nine taxa in the same function.

As previously assessed, *Asinibacterium* sp. has been described for dehalogenation capacities^55,57,59^ Even though their role cannot be excluded, all the other taxa recovered as harbored by the AN10-11Pos microbiome, were recorded at levels under the selected threshold of representation of the microbial load. *Dongia* sp., at the best of our knowledge has been described as negatively correlated with most organochlorine pesticides (OCPs)^61^. As far as we know any data have been previously reported for the *Stenotrophobacter* and *Nocardia* sps. and the transformation of any of the HCH isomers. *Sphimgomonas* sp. has been reported as capable to transform the β isomer ^36^. Moreover, Actinobacteriota and Chloroflexi have been reported as composing the microbial community of historical contaminated sites^60^.

Among the taxa that might be also involved in the highest efficiency of AN10-11Pos in β isomer depletion, two strains of the Hyphomicrobiaceae family, that characterize both the soil of origin and the microbiome of interest cannot be excluded. Hyphomicrobiaceae has been isolated from soils contaminated with HCH, although specific degradative abilities for individual isomers have not been identified^62^. A *Nocardioides* sp. taxa has also been recorded both in the AN10-11 Pos and in the soil of origin and the *Nacardioides* have been recorded as present in microbiomes responsible for the transformation of pesticides and in particular the β-HCH isomer^61^.

The co-metabolic enrichment conditions led to the enrichment of efficient microbiomes in depleting HCH, however, the most significant effect of glucose is the improvement of the microbiome β isomer transformation capacity. This metabolic trait that has been reported as possibly related to a multiple number of *lin*A gene in the genome of the single strain^62^. The genomic organization and stability of the *lin* genes have been also described as associated to the genomic organization ad copy number of the *IS1000* that has been reported as shaping the *lin* genes organization, due to the observed adjacent arrangement on the genome^62^. This insertion sequences are involved not only in the organization of the genome of the individual strain but also in the horizontal transmission of genetic information. In the context of the present study, the soil showing the highest bacterial community evenness might be reasonably associated to a wider distribution among different bacterial species of HCH transforming *lin* genes, favoring not necessarily richness and biodiversity of the bacterial community but the distribution of catabolic features among all the bacterial species that mutually resist to the toxicity of HCH contamination. The soil showing the highest evenness in bacterial diversity was associated to the lowest level of contamination. The two characteristics might be interdependent since at lowest level of contamination might correspond the lowest toxicity of the matrix and the higher evenness. However, the highest microbiome biodiversity was here demonstrated as mandatory for more favorable kinetics of transformation of the β isomers (AN10-11Pos *versus* AN7-8Pos). This assessment might suggest that the lowest level of contamination in a soil sample deriving from the same contaminated site, might be associated to an efficient exploitation of horizontal transmission of the catabolic activity among different bacterial taxa, with the consequent higher rate in HCH depletion in the soil and a higher evenness of the bacterial community responsible for HCH depletion. The assessment is suggested by the here observed possible stabilization of the microbiome basic metabolisms in presence of glucose that might have promoted the genomic arrangement of the *lin* genes within the microbiome’s metagenome, including the involvement of events of horizontal gene transfer and the increment of the biodiversity of bacteria capable to transform the HCH isomers. If this hypothesis is true, then it is important to consider the evenness of microbial communities colonizing matrices contaminated by HCH, when searching for a source to enrich microbiomes capable of degrading the complex mixture of characterizing isomers. Biodiversity indices could be important parameters to evaluate the potential natural attenuation capacity of a HCH contaminated matrix. In the near future, the adoption of a whole genome sequencing approach for the microbiomes under study will shed light on this hypothesis by annotating the *lin* genes and flanking insertion sequences in the genomes of the bacterial species composing the microbiomes.

## 5. Conclusions

Assessing that bioaugmentation is mandatory to accelerate the depletion of contaminants in the environment, and that microbiomes are essential effectors of environmental contamination degradation processes, it is necessary to redesign the approach for enriching candidates for bioaugmentation interventions. The present work has demonstrated that the biodiversity, being responsible for the richness of the metabolic and catabolic pathways of the reference microbiome, is essential in providing candidates with an elicited efficiency in the degradation of the most recalcitrant β-HCH isomer. A co-metabolic strategy of adaptation and enrichment, for toxic contaminants not providing sufficient reducing power to the microbiome of interest, such as HCH, resulted to be mandatory to sustain the above-mentioned biodiversity. An important role has been assessed for *Chryseobacterium* and *Asinibacterium* sps. in the depletion capacity of the β isomer.

## Acknowledgments

The research has been supported by the Ministry of University and Research (MUR) as part of the PON 2014 2020 Research and Innovation resources-Green/Innovation Action – DM MUR 1061/2022 and by Teseco Bonfiche co-financing the PhD fellowship of Giacomo Bernabei

## Authorship contribution statement

Giacomo Bernabei: Writing – original draft, Investigation, Methodology, Formal analysis, Conceptualization. Giampiero De Simone: Writing – original draft, Investigation, Formal analysis, Data curation. Simone Becarelli: Methodology, Formal analysis, Data curation. Riccardo Di Mambro: Writing – review & editing, Conceptualisation. Alessandro Gentini: Conceptualization. Simona Di Gregorio: Writing – review & editing, Supervision, Formal analysis, Conceptualization.

